# Multi-modal, multi-species, and multi-task latent-space model for decoding level of consciousness

**DOI:** 10.64898/2025.12.03.692227

**Authors:** Julia H. Wang, Méliya El Fakiri, Jake Swann, Elena Grazia Gado, Scott J.X. Cheng, Jordy Tasserie, Rongchen Huang, Jia Liu, Jongha Lee, Tianyang Ye, Paul Le Floch, Oliver Armitage

## Abstract

Assessing the degree and characteristics of consciousness is central to caring for patients with Disorders of Consciousness (DoC), yet current standard of care is a bedside questionnaire. Using a laminar neural probe we introduce a self-supervised, multi-modal representation learning approach that constructs an interpretable 2-D latent space of brain state from continuous neural recordings. We jointly encode local field potentials (LFPs), single-unit firing rates, and a movement proxy into a time-aware autoregressive VAE (TN-VAE). In rats, pigs and humans undergoing controlled depth anesthesia experiments, the learned latent trajectories nonlinearly, but smoothly, separate four expert-defined states (awake, light, moderate, deep) at 2*s* resolution, exceeding the temporal granularity of behavioral scoring (15*m*). The latent space supports linear readout of both coarse state and individual behavioral components, and its axes align with known physiology: delta/alpha power differentiates unconscious sub-states, while gamma and unit firing distinguish wakefulness. To enable cross-species use and incomplete modality sets, we add lightweight, species-specific stitching layers. This model pretrained on tri-modal rat data and fine-tuned with unimodal (LFP) pig data and unimodal human intraoperative data, successfully separates awake versus anesthetized states in new unseen human subjects, demonstrating zero-shot multi-subject transfer without per-patient calibration. The model further generalizes in a multi-task manner to predict the results of additional stimuli. These results highlight a path toward a foundation model for DoC that generalizes across sessions, subjects, species, and stimuli to enable scalable, continuous brain-state monitoring and clinical decision support.

## 1 Introduction

Disorders of Consciousness (DoC), including coma, unresponsive wakefulness syndrome, minimally conscious states, and covert consciousness, represent some of the most challenging conditions in clinical neurology. These conditions, typically following brain injury or illness, are characterized by impaired arousal and awareness—the fundamental abilities to remain awake and interact with one’s surroundings and self. While accurate diagnosis and prognosis are critical for providing early and personalized treatment, current standard-of-care methods rely on bedside assessments such as the Coma Recovery Scale-Revised (CRS-R) and Glasgow Coma Scale (GCS), yield misdiagnosis rates exceeding 40% (Andrews et al. [1996], Schnakers et al. [2009], Childs et al. [1993]), indicating significant need for technologies that can more accurately assess consciousness levels to guide therapeutic interventions and improve patient outcomes.

Brain-Computer Interfaces (BCIs) represent a promising solution for DoC patients who cannot respond to conventional assessments. For clinical deployment, these systems require zero-shot decoding capabilities, as they must function without patient-specific training or calibration. Neural foundation models provide a promising avenue for eventual clinical BCI applications as well as further scientific understanding of neural mechanisms underlying DoC (Wang et al. [2024], Ye et al. [2023], Kaifosh et al. [2025], Schneider et al. [2023]). Such foundation models should capture heterogeneity of modalities, sessions, and species as well as be useful in various downstream tasks Azabou et al. [2023]. To this end we employed anesthesia—a well-established, reversible model of consciousness transitions—to develop and validate a neural foundation model (Childs et al. [1993], Schnakers et al. [2009]).

Previous work in modeling consciousness has predominantly relied on supervised learning methods applied to unimodal EEG-derived features, requiring ground-truth signals such as behavioral responsiveness or anesthetic concentration that can be subjective or unreliable (Li et al. [2020], Afshar et al. [2021], Sun et al. [2019]), and may lack generalization across subjects (Li et al. [2020], Purdon et al. [2015]). Additionally, preliminary studies have demonstrated the potential benefits of integrating signals from different physiological systems in both clinical diagnosis and scientific understanding of underlying mechanisms for DoC (He et al. [2023], Brown et al. [2018])

To address these limitations, we developed a self-supervised deep representation learning model that integrates multimodal signals to create an interpretable latent space of consciousness depth. Using soft neural probes for stable single-neuron populations, implanted in both rodents and humans, we recorded LFPs, spiking activity, and movement signals. Our model predicts behavioral correlates of consciousness and anesthetic concentration while relating to existing clinical diagnostic methods for DoC, offering a foundation for improved consciousness assessment in clinical settings.

## 2 Methods

### 2.1 Depth of anesthesia experiments

We recorded neural signal from the brain of a rat subject implanted with Fleuron™ probes targeting the right hemisphere, spanning the somatosensory cortex and hippocampus (Fig. 1b, Appendix A.1) (Lee et al. [2025]). These neural signals were then used to extract both LFP signal, firing rates of individual neurons, and a proxy of movement (Appendices A.4 - A.6). We recorded during five depth of anesthesia experiments, each on separate days over multiple weeks, in which we injected the subject with various doses of a ketamine/xylazine cocktail. As the subject transitioned to deep anesthesia and then back to wake through several levels of consciousness, we continuously monitored state through a behavioral scoring system (Fig. 1a). Briefly, we assessed the breathing rate, exploration of area, spontaneous movement, level of balancing, and toe pinch response of the subject throughout the experiment approximately every 30 seconds. These scores are combined to give an overall behavior score and assigned to one of 4 broad anesthesia states: awake, light, moderate, and deep (Appendix A.2). We repeated this experiment in a pig subject over two sessions. We further recorded neural signal from 3 human subjects under different states of propofol anesthesia during functional neurosurgery, while also playing an audio oddball, local-global paradigm, to test consciousness (Appendix A.3, A.11).

**Figure 1.**
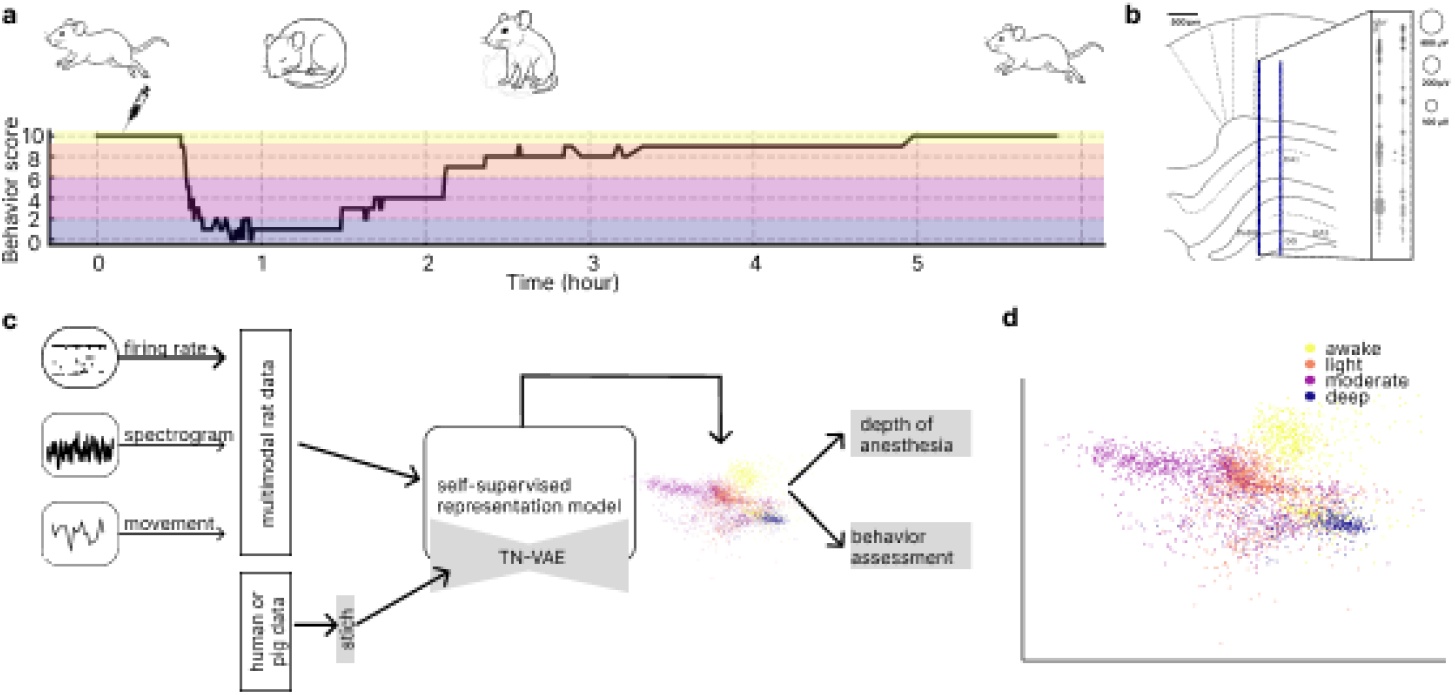
**a)** Timeline of a representative recording session spanning wakefulness, depths of anesthesia, and recovery. Behavioral state was scored throughout (Appendix A.2). The syringe marks the time of the ketamine/xylazine (K/X) injection. **b)** Left: Rat brain atlas showing the location of implanted probe across the cortex and hippocampus. Right: Distributions of amplitude and depth (centroid) for detected units from the same session. **c)** Schematic of multimodal self-supervised representation model. Firing rates of individual neurons LFP wavelets, and estimated movement are used to create a multimodal feature representation of any point in time. This feature is passed to an auto-regressive variational autoencoder known as TN-VAE. The 2-D latent space of the autoencoder is then used to predict anesthetic concentration and behavioral assessments, which are not supervised or input signals to the model. The model handles multi-species data with different modalities through species-specific stitching layers. **d)** Latent space from the unseen test session separates anesthesia states.

### 2.2 Self-supervised multi-modal representation model

For each 2 seconds of data, we construct a multi-modal feature representation by combining LFP spectral information, firing rates of spike-sorted neurons, and estimated movement data. This multi-modal representation is passed into a self-supervised representation model, TN-VAE (Fig 1c) (Wang et al. [2023]) to construct an interpretable 2-D latent space (Fig. 1d). The TN-VAE is an autoregressive variational autoencoder that seeks to promote smoothness in representations over time, and has been shown previously to be effective in separating sleep brain states from neural data. The model is trained with a dataset spanning over 100 channels and containing data from multiple recording sessions across multiple weeks. Importantly the model is able to integrate data from these sessions that are on separate days more than a month apart and constructs a continuous latent space without separating per session. In order to account for multi-species data, in which each species might have arbitrary missing or extra modalities, we developed species-specific stitching layers that can transform the species-specific modalities to the latent space of the base model (Fig 1c) (Turaga et al. [2013], Ye et al. [2023]). These stitching layers can be used to train either a multi-species model, or in a transfer learning regime with limited amounts of data.

## 3 Results

### 3.1 Self-supervised model separates states of consciousness

After training, we applied our model to channels from an unseen test session from the same rat spanning multiple states of consciousness. Our model effectively separates the 4 states of anesthesia (Fig. 1e, Fig. 2f, Appendix A.9). The separation of states by a non-linear deep learning model exceeds that of a principal components baseline model based on linear decodability of the latent space to behavior scores (Fig. 4a,b). Furthermore, the state score was assessed at a 30 second time resolution, whereas our model is able to distinguish state at a 2 second resolution, explaining some mismatch between our representation space and behavior score. The model traverses the latent space non-linearly but smoothly through time, returning to the initial position in the latent space, as the subject starts and ends in the wake state (Fig 1a,b). We additionally estimated anesthetic concentration in the subject based on the timing of the ketamine/xylazine cocktail administration and known pharmacokinetic constants (Appendix A.9). We see that the anesthetic concentration does not track linearly with the behavior score; both manual observation of consciousness level through behavior and neural signal can contain finer-grained information than simple anesthetic administration calculations (Fig 1c). Through examining how sub-behavior scores tile the latent space, we also find that while these behavior signals are somewhat separated in latent space, no single behavior is sufficient in determining anesthesia state. Exploration of area is important in separating awake from other states, balancing for distinguishing between light and moderate, and toe pinch response differentiates the deepest states (Fig. 1d). This is further illustrated by the linear decodability of individual behavioral signals vs broad behavior state from the latent space (Fig. 1f).

**Figure 2.**
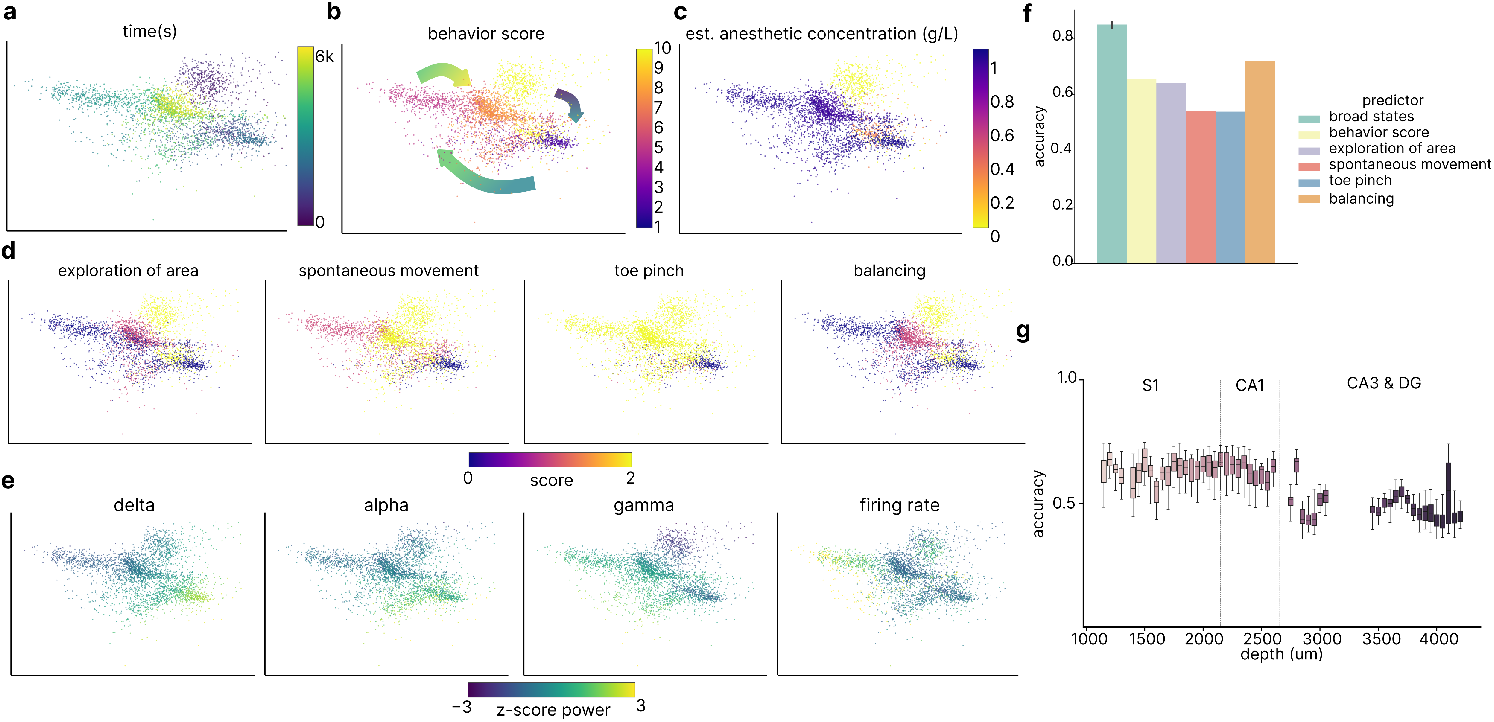
**a)** Latent space of example channel from the unseen test session colored by time in the session **b)** latent space colored by overall behavior score, with arrows showing trajectory over time. **c)** Example latent space colored by estimated anesthetic concentration. **d)** Example latent state colored by behavioral score for (left to right) exploration of area, spontaneous movement, balancing, and toe pinch. **e)** Example latent space colored by different features, such as average power in differently LFP frequency bands and single neuron firing rates. **f)** Linear decodability of the four broad states, overall behavior score, and individual behavior scores for the unseen test session. Error bars are over train/test splits and decoder initializations. **g)** Linear decodability of broad states (awake, light, moderate, deep) across depths of the probe. Whiskers are over channels of the same depth and multiple classifier instantiations.

### 3.2 Interpretability of latent space

Next we interpreted which features were most important in creating our latent space. In accordance with previous understanding of LFP frequency bands in anesthetic states, we find that alpha and delta are important for separating between deep, moderate, and light; while gamma is useful in separating wake from unconscious (Fig. 2e). We find that the firing rate of individual neurons shows additional variation within the 4 broad states beyond that of the LFP frequency bands, which may indicate usefulness in creating a continuum of states in the latent space and will be an interesting avenue for further study (Fig. 2e). We find that anesthesia states are decodable across the regions (Fig. 2g).

### 3.3 Self-supervised model learns across species and contains multi-task information

We next fine-tuned our model pretrained on trimodal data from rat with unimodal (LFP) human data. We trained with only 18 minutes of data that encompassed unconscious and awake states in one human subject, and found that our model was able to separate states recorded from two unseen human subjects(Fig. 3a). The label of the unconscious to conscious transition was based on anesthesiologist assessment of the patient and is therefore not exact to a second resolution and does not capture the transition of regaining consciousness. Nevertheless, the model is able to separate these distinct states, and could be used to further understand this transition across channels or regions. In addition to decoding level of consciousness from passive data, our model demonstrates multi-task performance on this dataset by capturing the neural responses to applied audio stimuli whose neural responses are known to depend on state of consciousness (Fig. 3b,c, Appendix A.11). Briefly, when processing the neural response to audio tones the model captures in the latent space whether the response is indicative of consciousness or unconsciousness. The subject B latent space includes responses to audio indicating local pattern deviation processing, the subject C subset of the latent space includes two distinct regions of responses to audio indicating local and global pattern deviation processing. Lastly, we also fine-tuned the model from bimodal (LFP and movement) data from a pig subject. The model was trained on data from one session and applied to another. In both sessions, the pig spanned between light and deep anesthesia states, and the latent space is able to separate these states from each other (Fig. 3c).

**Figure 3.**
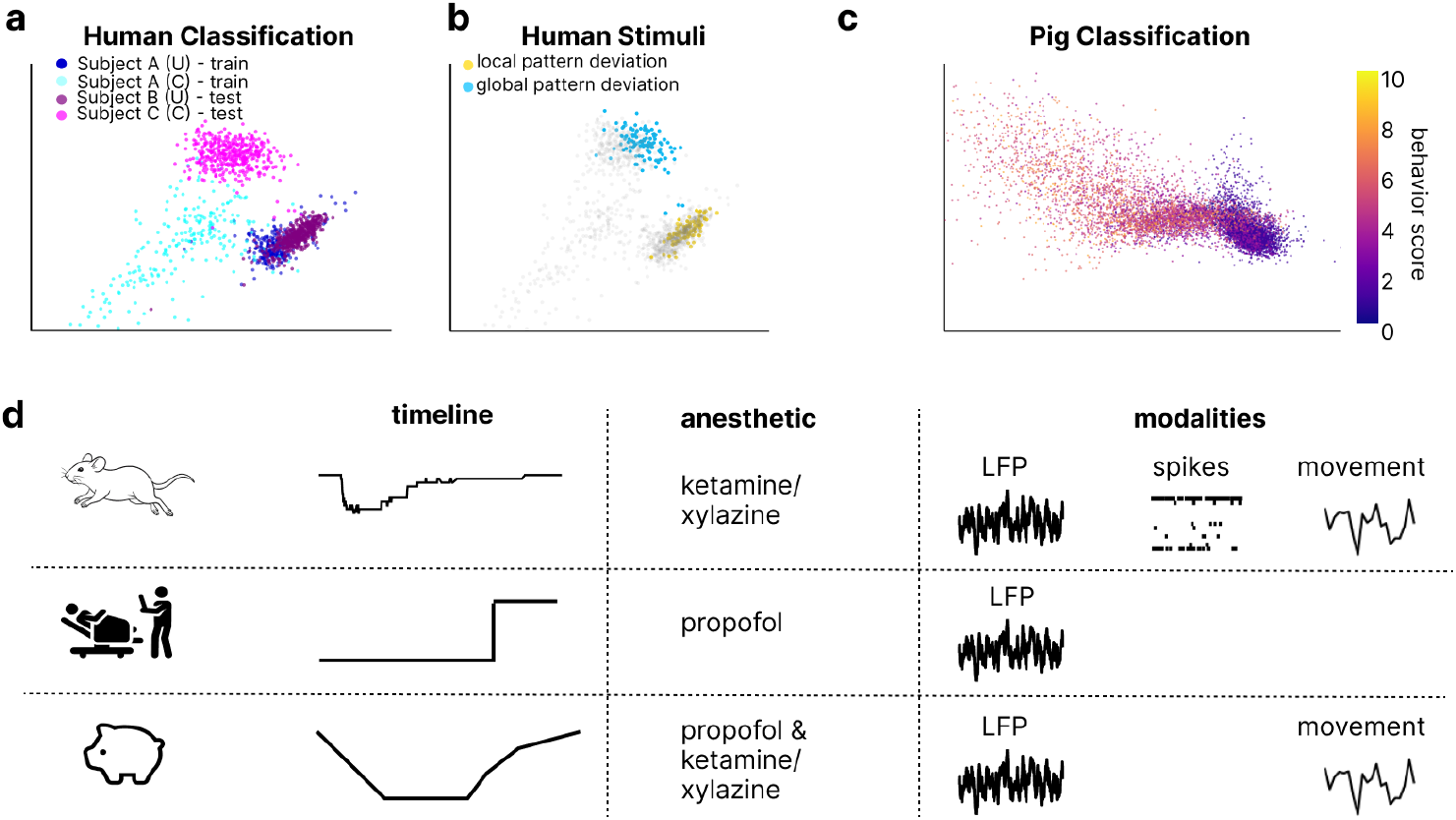
**a)** Latent space of example channel from a model fine-tuned on human data from one subject that transitioned between unconscious and conscious states. This model is then applied to two unseen subjects, one unconscious and one conscious. **b)** Time points in which audio stimulus tones played as colored by measuring local response or global response. While local response metric is consistent across human subjects irrespective of consciousness state, global response metric determines consciousness states. **c**) Latent space of an example channel fine-tuned on pig data from one subject on an unseen test session. **d)** Schematic of data from a variety of species such as rat, human, and pig. Additionally different anesthetics are given resulting in different consciousness state timelines, and different data modalities recorded, such as LFP, spiking data, and auxillary behavioral signals.

**Figure 4.**
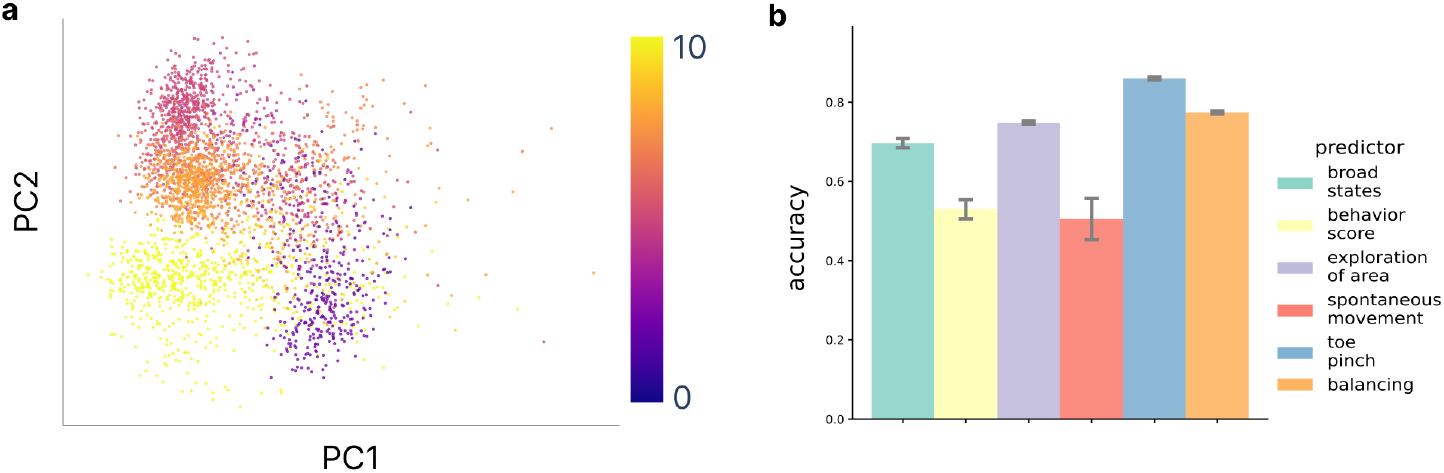
**a)** Principal component space of the test session of rat anesthetic states colored by behavior score. The training and test set is the same as in Figure 2. **b)** Linear decodability of the four broad states, overall behavior score, and individual behavior scores from the first 2 PCs.

## 4 Limitations

Herein we present results of our multi-session, multi-modal multi-species model that can generalize across these cases. The rat data is extensive in time but limited in that it is only from a single subject. The human data is multi-subject, but due to the constraints of the surgery paradigm, totals only 100*min* in length and as such the class separation, while successful, could be improved. We also find that our model is sensitive to artifacts and distributional effects across channels, sessions, and subjects, which can be solved by z-scoring, but remains an area of exploration. In future studies, we hope to incorporate other external modes of behavior such as behavioral videos, heart rate, breathing rate, and more. The flexibility of the model allows these signals to be integrated as long as they are presented continuously in time. Lastly, our model creates a latent space per channel, rather than a latent space that represents activity combined from all channels. This allows our model more flexibility in transferring to new subjects or species that may have different number of channels, but our future work will extend to creating multi-channel representations.

## 5 Discussion

This work lays the groundwork for a brain foundation model to improve diagnostic accuracy and prognostic assessment in patients with disorders of consciousness (DoC). By generalizing across subjects and modalities with minimal training data, the model shows promise for diagnosing new patients without individual calibration. Continuous monitoring of multimodal neural signatures may reduce diagnostic uncertainty, ease the burden on clinicians and caregivers, and support targeted therapies. We expect the multi-modal and multi-species nature of the model will improve accuracy and task output, allowing for few-shot fine tuning on new subjects. While anesthesia offers a controlled model of consciousness transitions, it does not fully capture DoC’s complexity, highlighting the need for models trained on diverse brain states to guide further clinical assessment. Current computational models often overlook brain state as a core factor in cognition, treating it as variability; incorporating explicit brain state modeling—especially using both single-neuron activity and broader network dynamics—could enhance predictions across cognitive domains like language and motor control.

## A Technical Appendices and Supplementary Material

### A.1 Rat surgery and recordings

All rat experiments conducted in this study were approved by the Mispro Institutional Animal Care and Use Committee under protocol AXO-2022-01. 11 week old Sprague Dawley rat implanted with Fleuron™ probes targeting the right hemisphere (AP -3.5 mm, ML +1.8 mm, DV 4.2mm) was used in this study (Lee et al., 2025). Electrophysiological signals were amplified and digitized with a custom recording system integrated with RHX Data Acquisition Software (INTAN Technologies). A custom-printed circuit board (PCB) connected the implanted probes to the recording system. Neural signals were acquired at 20 kHz sampling frequency with a 60 Hz notch filter. Electrode impedance was measured at 1 kHz using the same recording setup. A detailed overview of Fleuron™ neural probes and their implantation in rats and humans can be found in Lee et al. [2025].

### A.2 Rat depth of anesthesia experiment and behavior scoring

Depth of anesthesia recordings were performed by administering ketamine/xylazine cocktail to create a wake to anesthesia and returning to awake state session. We developed a behavioral scoring system to characterize rat behavior across wakefulness, anesthesia, and recovery, adapted from clinical arousal scales in humans and nonhuman primates (Uhrig et al. [2016], Tasserie et al. [2022]) and informed by our pilot observations in anesthetized rats, while incorporating behavioral observations made by our investigator team in rats that underwent surgical anesthesia. A trained experimenter scored the behavior from following five domains and summed them to yield a composite score (higher = more awake): environmental exploration (0 = none; 1 = limited sniffing/locomotion within the arena; 2 = active investigation, e.g., head/ear orienting to sounds), spontaneous movement (0 = none; 1 = eye blink and/or whisker twitch; 2 = limb or whole-body movement), respiratory rate (0 = very slow, *≤* 60 bpm; 1 = 60 *≥ −* 90 bpm; 2 = *≥* 90 bpm), response to toe pinch (0 = none; 1 = slight withdrawal; 2 = clear withdrawal/strong movement), postural control/balance (0 = no movement; 1 = unsteady; 2 = coordinated walking).

### A.3 Pig surgery and depth of anesthesia experiment

All pig experiments conducted in this study were approved by the IPST Animal Care and Use Committee. Landrace pigs around 50kg were implanted with Fleuron™ probes targeting the right rostrum gyrus and used in this study. A ketamine/xylazine mix was used to induce anesthetic state in two sessions across multiple weeks. A trained experimenter scored the behavior from following five domains and summed them to yield a composite score (higher = more awake): jaw tone, spontaneous movement, breathing rate, palpebral reflex, and balancing.

### A.4 Human neural recordings

Human neural signal recording was approved by the Ethics Committee of The Panama Clinic and is registered at ClinicalTrials.gov under study number NCT06673264. We further recorded neural signal using the same Fleuron™ probes in 5 humans undergoing planned tumor resection surgeries. Two subjects were unconscious for the duration of the neural recording under a propofol anesthesia, one subject was fully conscious for the duration of the neural recording while undergoing functional neurosurgery, and two subjects started the neural recording unconscious and propofol anasthesia was withdrawn during neural recording and the subject gained consciousness. Subjects state of consciousness was assessed by the attending anesthesiologist.

### A.5 Data preprocessing: LFP wavelet calculation

To obtain the LFP band of the neural signal, we downsampled the neural signal to 500 Hz and bandpass filtered to between frequencies of 0.5 Hz and 300 Hz. We also applied notch filters at 60, 70, 120, and 156 Hz to remove electrical noise artifacts. We then calculated complex morlet wavelets of 50 scales corresponding to a log scale between 0.5 Hz and 150 Hz frequencies.

### A.6 Data preprocessing: spike sorting and firing rate estimation

Data was highpass filtered at 300Hz, and spike sorting was performed using Kilosort 4 (Pachitariu et al. [2024]). The initial automated sorting results were then curated using in one of two ways. At first, curation was done manually in the Phy GUI (https://github.com/kwikteam/phy). Once a number of recordings had been performed and curated thus, a machine learning classifier was trained on existing curated data to perform curation on future spike-sorting results, using the UnitRefine (Jain et al. [2025]) implementation in the Python package SpikeInterface (Buccino et al. [2020]). Subsequent curation was performed using this trained classifier to assign quality labels to each cluster based on a range of computed quality metrics. Units were assigned to depth bins based on their estimated position along the probe, as determined by template extremum channels and probe geometry. The probe was divided into contiguous 200*µm* depth bins. For each depth bin, all units located within its boundaries were grouped, and their spike counts were averaged to produce a firing rate time series per depth bin. Each channel was then associated with one or more depth bins, depending on the spatial distribution of units.

### A.7 Data preprocessing: movement signal estimation

We adapted a previously described approach to compute an estimation of the the electromyographic signal (EMG) originating from the snout, mouth, and shoulder muscles of the rat. To obtain this estimate, we computed zero time-lag Pearson correlation coefficients between high-frequency components of the intracranial recording. For each recording session, channel pairs were selected on opposite sides of the shank at 400*µm* spacing, sampled across four depths (superficial, 33%, 66%, and maximum depth). For each pair, signals were filtered in the 300–600 Hz band with transition bands extending from 275 to 625 Hz. Zero-lag Pearson correlation was then computed. and correlations were then averaged across pairs to yield an estimated EMG score with 1 Hz resolution.

### A.8 Dataset

We used a combination of depth of anesthesia sessions and other recordings which were performed under freely moving conditions. In total, 11 continuous recordings were collected over 6 weeks, resulting in a total of approximately 13.5 hours of data. We used 10 recordings in our training set, and held out 1 depth of anesthesia recording session for test. Each datapoint in our dataset corresponds to a 2 second window of time, and contains information including the LFP wavelets from a single channel, the total firing rate of neurons for that channel bin, and the movement data at that time, which is shared across all channels. We excluded noisy channels through a median absolute deviation noise estimation, and we removed artifact time periods throughout the experiment based on when the experimentalist manipulated the mouse, resulting in noisy signal. We created training and test datasets combining data across channels and sessions. We z-scored the features per channel and session to remove batch effects.

### A.9 Model architecture and training details

We adapt a previously described autoregressive VAE known as the TN-VAE Wang et al. [2023]. Briefly the model aims to predict the representation of the next point in time rather than reconstruct the original point. The autoencoder creates a 2-D latent space through a bottleneck layer. We input our multimodal representation (LFP, spikes, movement) into the model. We performed a hyperparameter sweep across several learning rates, batch sizes, KL divergence weights, and epochs. The training set was split randomly into a training(80%) and validation(20%) set for each model initialization. The final model was selected through the neighbor loss as described previously, and the final hyperparameters were as such: Adam optimizer with lr 1e-4, batch size 10000, kl weight 1e-4, epochs 200.

### A.10 Anesthetic concentration estimation

A relative anesthetic concentration was estimated using pharmacokinetic constants of ketamine and xylazine as described in Veilleux-Lemieux et al. [2013].

### A.11 Linear decodability of the latent space

We described the linear decodability of different behavioral states within a session. For the broad behavior states of (awake, light, moderate, deep), we trained a support vector classifier and for other behavior scores, we trained a ridge linear regressor. Mean accuracies and error bars (95% confidence interval) were taken over 100 trials with different train/test splits and initializations.

### A.12 Auditory stimulus response

We performed an auditory local-global deviant paradigm (Bekinschtein et al. [2009], Chennu et al. [2013]) for the 3 human subjects. We then defined a response metric as the negative correlation coefficient between the average LFP power in response to rare and frequent trials (global) or standard and deviant trials (local).

### A.13 Compute resources

Models were trained with an internal cluster that contained several A5000 GPUs.

